# Origin of Slow Spontaneous Resting-State Neuronal Fluctuations in Brain Networks

**DOI:** 10.1101/186098

**Authors:** Giri P. Krishnan, Oscar C. González, Maxim Bazhenov

## Abstract

Resting or baseline state low frequency (0.01-0.2 Hz) brain activity has been observed in fMRI, EEG and LFP recordings. These fluctuations were found to be correlated across brain regions, and are thought to reflect neuronal activity fluctuations between functionally connected areas of the brain. However, the origin of these infra-slow fluctuations remains unknown. Here, using a detailed computational model of the brain network, we show that spontaneous infra-slow (< 0.05 Hz) fluctuations could originate due to the ion concentration dynamics. The computational model implemented dynamics for intra and extracellular K^+^ and Na^+^ and intracellular Cl^-^ ions, Na^+^/K^+^ exchange pump, and KCC2 co-transporter. In the network model representing resting awake-like brain state, we observed slow fluctuations in the extracellular K^+^ concentration, Na^+^/K^+^ pump activation, firing rate of neurons and local field potentials. Holding K^+^ concentration constant prevented generation of these fluctuations. The amplitude and peak frequency of this activity were modulated by Na^+^/K^+^ pump, AMPA/GABA synaptic currents and glial properties. Further, in a large-scale network with long-range connections based on CoCoMac connectivity data, the infra-slow fluctuations became synchronized among remote clusters similar to the resting-state networks observed *in vivo*. Overall, our study proposes that ion concentration dynamics mediated by neuronal and glial activity may contribute to the generation of very slow spontaneous fluctuations of brain activity that are observed as the resting-state fluctuations in fMRI and EEG recordings.

## Introduction

Resting-state or spontaneous background fluctuations, in the frequency range of 0.01-0.2 Hz (1-18), have been reported by a wide range of neuroimaging methods, including electrophysiological, optical, EEG and fMRI (4, 6, 7, 10, 16, 19). The spontaneous resting-state activity in fMRI signal is a robust phenomenon that has been widely used to evaluate brain network properties, from determining functional connectivity during cognitive tasks to identifying altered functional connectivity in various conscious and disease states (1, 2, 4, 6, 7, 17, 20, 21). The resting-state activity across wide brain regions forms functional networks, such as the default-mode network (DMN), that vary with brain state and type of cognitive activity (1, 2, 4, 7, 17, 20). Several neurological and psychiatric disorders, such as epilepsy and schizophrenia, have been shown to correlate with altered resting-state fluctuations and functional connectivity (1, 2, 6, 20-24). Although there is growing interest in understanding resting-state fluctuations, the underlying neural mechanisms by which these oscillations arise remain unknown.

Previous experimental work showed that infra-slow fluctuations in the local field potential (LFP) gamma power, neuronal firing rate, and slow cortical potentials (SCP) exhibit a correlational relationship with resting-state fMRI blood-oxygen-level dependent (BOLD) fluctuations (9, 10, 12, 15-17). Further, the underlying structural connectivity of brain network was shown to shape the functional connectivity estimated from resting-state activity (8, 11, 25, 26). Finally, various computational studies, using population level models, suggested the role of intrinsic noise, coupling strengths, conduction velocities, and underlying structural connectivity in the generation of resting-state fluctuations (5, 11, 25). Nevertheless, the mechanistic origin of the infra-slow time scale based on activity of individual neurons and their networks remains to be understood. Interestingly, experimental and computational studies suggest that the resting-state activity arises from switching between different activity levels in the localized brain regions. Changes in the ion concentrations have been suggested to modulate network activity (27-34) that could occur at a slow time scale. In this study, we tested the hypothesis that the infra-slow fluctuations of the ion concentrations may result in the resting-state fluctuations.

Extracellular potassium concentrations ([K^+^]_o_) have been shown to fluctuate during resting-state or background activity over a long time period (35). Recordings from anesthetized cat cortex have shown that the [K^+^]_o_ exhibits small amplitude (~ 0.5mM) fluctuations around a mean concentration, with the period of these fluctuations being about an hour or less (35). Additionally, extracellular potassium recordings in animal models of epilepsy have shown substantial [K^+^]_o_ fluctuations prior to and during bouts of seizure-like activity (32, 33, 36). A number of computational models suggested a prominent role of potassium concentration dynamics modulating neuronal excitability and synchrony (27-31, 34, 37). In this new work, we found that resting-state activity can arise from infra-slow fluctuations in the ion concentrations. Furthermore, we identified that the low amplitude fluctuations in the ion dynamics allow for the local and long-range synchronization among the distinct networks of neurons. These local fluctuations can generate correlated and anti-correlated activities between clusters of neurons through long-range feedforward excitation and inhibition, which reflect underlying network structural connectivity in agreement to the experimental observations of the resting-state activity.

## Results

The biophysical network model incorporated synaptically coupled excitatory (PY) and inhibitory (IN) neurons and implemented realistic dynamics of the major ion concentrations to provide *in vivo*-like conditions (28-31, 37). A “single cluster” network model possessed only local connectivity (5 neuron radius). Below we will fist explore dynamics of a single cluster network and then expand it to the case of several clusters connected with long-range synaptic connectivity.

In a network consisting of a single cluster of 50 neurons (Figure 1 A1), PY and IN population activity appeared random over a period of 800 sec simulation time (Figure 1 A2 top). Individual voltage traces from PY neurons within the network showed spontaneous random firing (Figure 1 A2 bottom). However, analysis of the mean firing rate of PY neurons revealed a very slow semi-periodic fluctuation (Figure 1B). Similar fluctuations were observed in the band-pass filtered mean PY membrane potential (Figure 1C). The slow fluctuations in the mean firing rate and membrane potential were also reflected in the slow fluctuations of the mean [K^+^]_o_ and [Na^+^]_i_ (Figure 1D green and red respectively), and the mean Na^+^/K^+^ pump current (Figure 1E). The [K^+^]o fluctuation was 0.1-0.2 mM in amplitude, while the [Na^+^]_i_ fluctuation was only ~0.05 mM in amplitude. Fourier transforms of the mean [K^+^]_o_, [Na^+^]_i_, Na^+^/K^+^ pump current, membrane potential, and network firing rate revealed a peak amplitude around ~0.02 Hz (Figure 1F).

**Figure 1.**
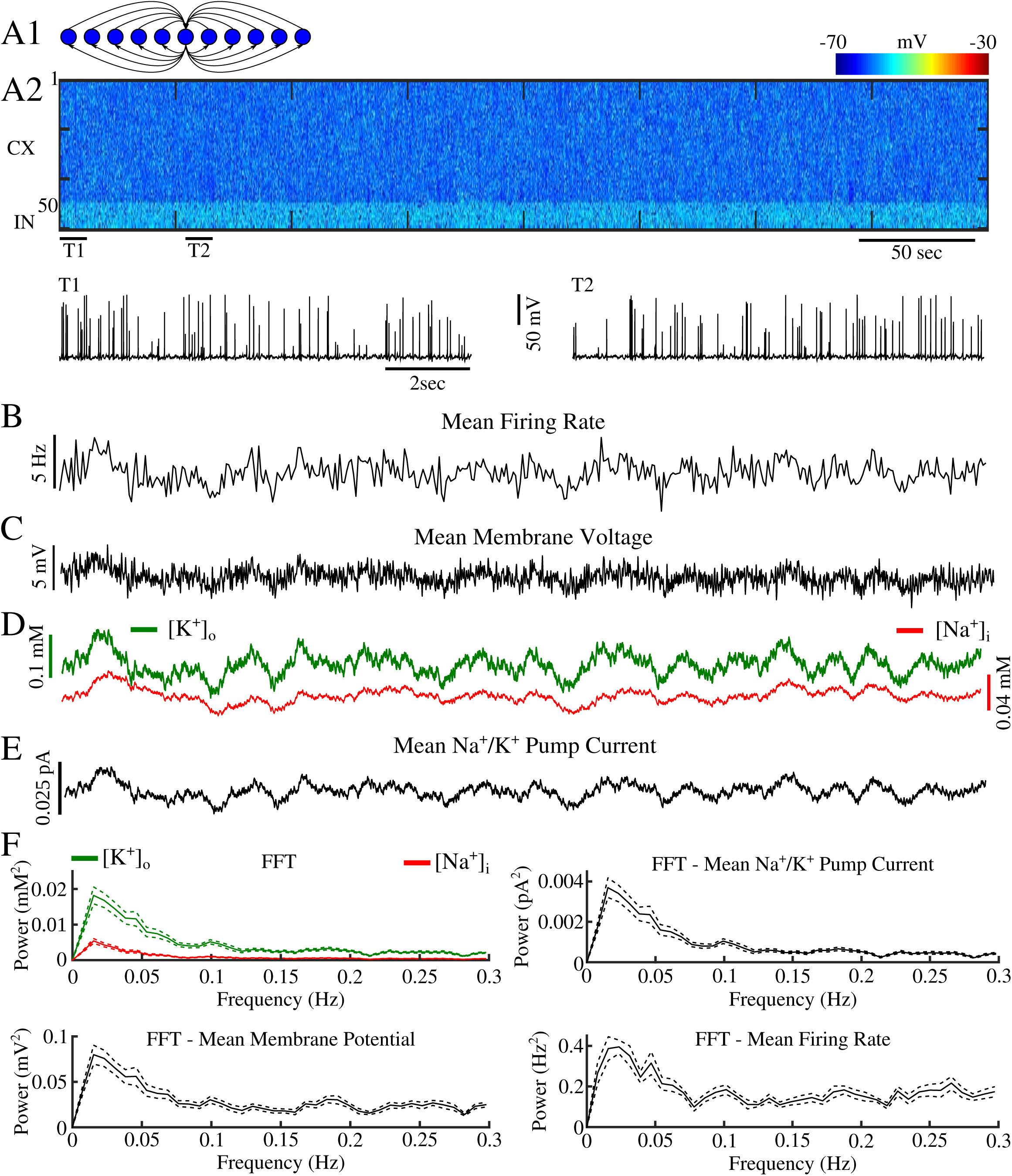
Resting-state network exhibits resting-state fluctuations. **A1,** Cartoon of basic network connectivity. **A2,** Spontaneous activity in a network of PY and IN neurons. T1 and T2 indicate time of zoomed in single cell traces from a single PY neuron. **B**-**E,** mean PY firing rate, mean filtered (0.001-0.1Hz) membrane potential, [K^+^]_o_ & [Na^+^]_i_, and Na^+^/K^+^ pump current, respectively. **F,** Fourier transforms of mean PY firing rate, membrane potential, [K^+^]_o_ & [Na^+^]_i_, and Na^+^/K^+^ pump current.

Since we observed very slow resting-state fluctuations in both mean [K^+^]_o_ and [Na^+^]_i_, we first tested the role of these ions in the generation of the resting-state fluctuations in our network model. Thus, we fixed the concentrations of these two ions independently. Holding the [K^+^]_o_ concentration constant resulted in a loss of the infra-slow oscillatory activity in the mean membrane potential of the PY neurons (Figure 2A top red). Fourier transform revealed disappearance of a peak at around 0.02 Hz and almost flat power spectrum (Figure 2B). In contrast, preventing the [Na^+^]_i_ fluctuation did not result in a loss of the slow membrane potential fluctuations (Figure 2B top & bottom). Unlike the condition with fixed [K^+^]_o_, the Fourier transform of the mean membrane potential in the network with constant [Na^+^]_i_ still showed a distinct peak around ~ 0.02 Hz. This suggested that fluctuations of [K^+^]_o_ play a major role in the generation of the resting-state activity, while [Na^+^]_i_ fluctuation may play a modulatory role.

**Figure 2.**
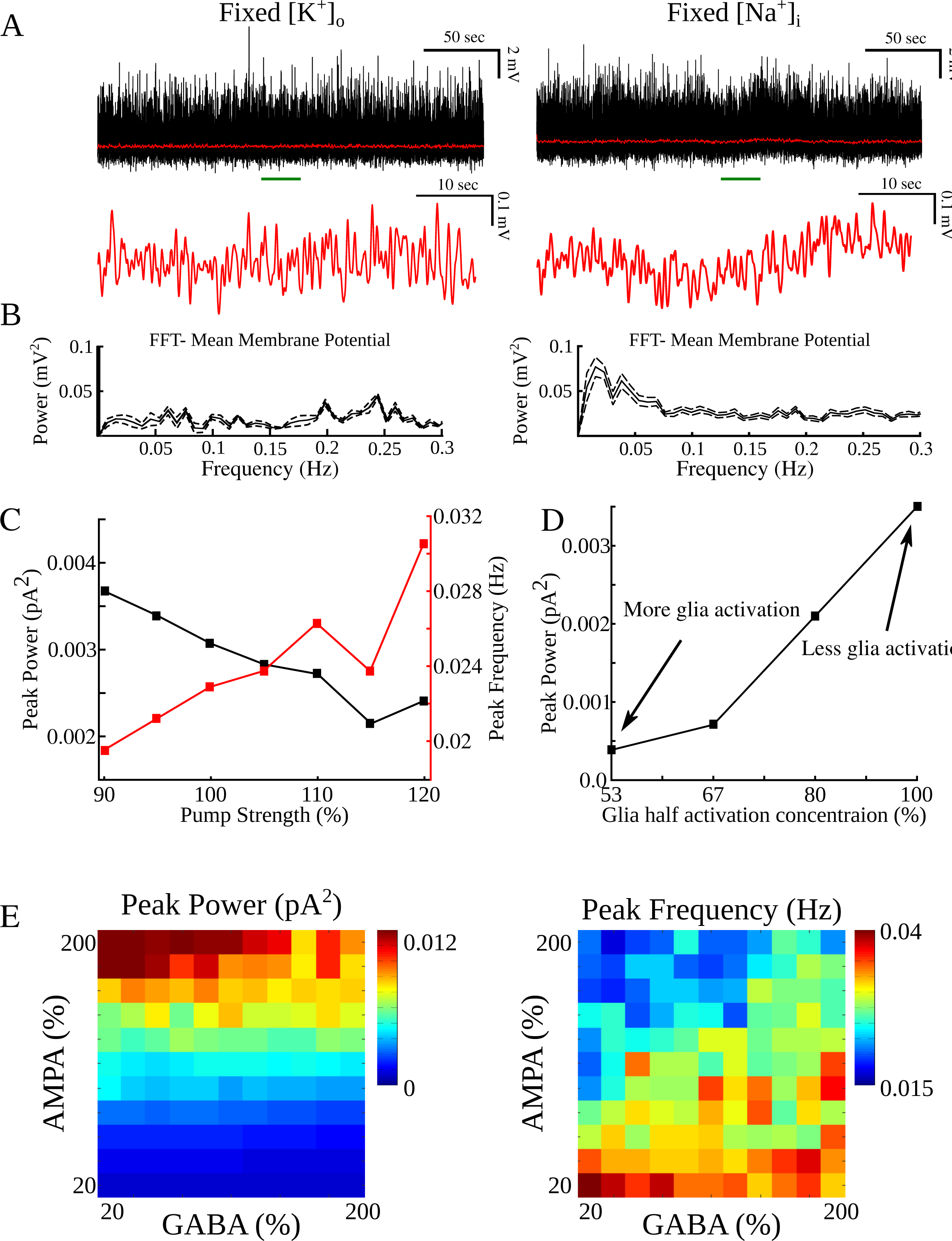
Ion concentration fluctuations, Na^+^/K^+^ pump, glial K^+^ buffering, and AMPA/GABA strength influence resting-state fluctuation properties. **A,** Mean membrane potentials (black) and mean filtered membrane potentials (red) for networks with either fixed [K^+^]_o_ or fixed [Na^+^]_i_ (left and right respectively). **B,** Corresponding Fourier transforms (FFT) for mean membrane potentials in A. **C,** FFT peak power and peak frequency as a function of Na^+^/K^+^ pump current strength (black and red, respectively). **D,** FFT peak power as a function of glia half activation K^+^ concentration. **E,** FFT peak power as a function of both AMPA and GABA connection strengths (left). FFT peak frequencies as a function of both AMPA and GABA connection strengths (right).

We next explored the role the Na^+^/K^+^ pump current, glia K^+^ buffering, and AMPA/GABA synaptic connections in affecting the properties of the resting-state fluctuations. Increasing the strength of the Na^+^/K^+^ pump resulted in decreased fluctuation amplitude of the pump current as revealed by the reduced peak in the Power spectrum (Figure 2C, black). This decrease in the fluctuation amplitude was accompanied by increase in the peak frequency (Figure 2C, red). For higher strength of the Na^+^/K^+^ pump, the relative changes of the extracellular K^+^ were reduced, leading to the smaller resting-state fluctuations. Similarly, decreasing the half activation concentration of [K^+^]_o_ in glia cells, that increased glia buffering, reduced the resting-state fluctuation amplitude (figure 2D). Thus, the slow dynamic processes controlling progressive accumulation of the ions and their removal determined the amplitude and the peak frequency of the slow fluctuations. Further, increasing the strength of the AMPA connections between PY neurons increased the amplitude of the resting-state fluctuations while shifted peak frequency to the lower values (figure 2E). GABA, on the other hand, had minimal impact on the amplitude of the resting-state fluctuations, but shifted the peak frequency to the higher values (figure 2E). Increase in the fluctuation amplitude due to increase in the AMPA connection strength suggests that recurrent synaptic excitation promoted faster and higher level buildup of the extracellular K^+^ concentration thus leading to stronger and faster resting-state fluctuations.

Does the network size affect generation of the resting-state activity? To answer this question, we varied the number of neurons in the network, but kept the same network configuration as in shown in Figure 1 A1. Again, the network contained only local connections (i.e. 5 neuron radius), and the ratio of PY to IN neurons was kept at 5:1. Smaller networks, 20–50 neurons, revealed large fluctuations in the mean Na^+^/K^+^ pump current (Figure 3A right) while the peak frequency remained the same around 0.02 Hz (Figure 3A left). In contrast, as the number of PY neurons in the network increased beyond 100 neurons, the amplitude of the mean (averaged over entire network) Na^+^/K^+^ pump current fluctuation decreased, which was revealed by the Power spectrum analysis (Figure 3A). The 500 neuron network revealed the smallest fluctuations and lowest power spectrum peak around 0.02 Hz. In order to get a better understanding of this phenomenon, we looked for the formation of local clusters within the 500 neuron network. We found that this network did in fact develop 5 local clusters of about 100 synchronized neurons each (Figure 3B). These local clusters formed spontaneously, and each cluster showed larger Na^+^/K^+^ pump current fluctuations than the pump current averaged over the entire network (Figure 3B). We also observed the recovery of the power spectrum peak amplitude when looking at the individual clusters as opposed to the entire network (Figure 3C). This suggests that local clusters can form spontaneously in large networks containing local network connectivity and dynamic ion concentrations, and that the resting-state fluctuations in these clusters can form complex phase relationships among them.

**Figure 3.**
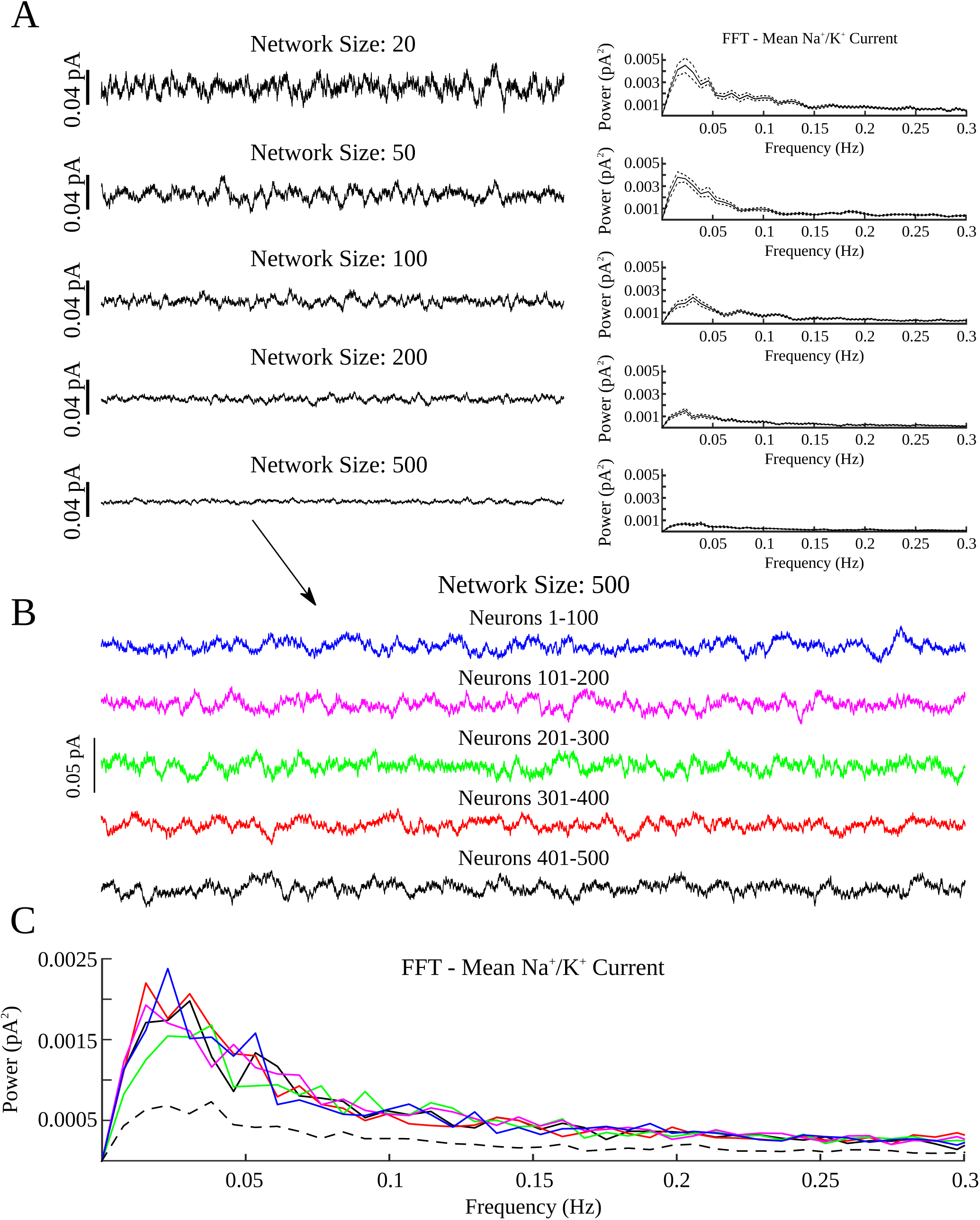
Effects of network size on resting-state fluctuation amplitude. **A,** Mean Na^+^/K^+^ pump currents for network so different sizes (left) and corresponding FFTs (right). **B,** Mean Na^+^/K^+^ pump currents from bin of 100 neurons comprising the network of 500 neurons in A. **C,** Corresponding FFTs for Na^+^/K^+^ pump currents (colored lines) in B, and FFT of Na^+^/K^+^ pump current from entire 500 neurons (dashed line).

Structures comprising functional networks, such as the ventromedial prefrontal cortex (vmPFC) and posterior cingulate cortex (PCC) of the default mode network, have been shown to have coherent resting-state fluctuations (1, 2, 7). Though fluctuations of the local ion concentrations, as proposed by our study, may underlie intrinsic fluctuations in these regions, the spatial separation of these regions makes sharing of the extracellular space between them unlikely. Therefore, these distinct regions should have independent dynamics of the local milieu of the ion-concentrations. On the other hand, many brain regions are known to be connected through long-range synaptic projections. Thus, we next tested whether our model could also generate coherent fluctuations between distinct clusters of neurons connected through long-range synaptic connections where each cluster has local synaptic connectivity and local extracellular ion concentration dynamics. As depicted in figure 4 A1, our new network configuration consisted of two clusters of 50 excitatory neurons and 10 inhibitory interneurons each. We kept the same connectivity scheme as the previous network configuration (Figure 1 A1) within each neuron cluster, but prevented ions from diffusing between the two clusters. Additionally, we added long-range sparse synaptic projections between two clusters through excitatory PY-PY connections. Again, the activity of the PY and IN neurons in either cluster appeared to be random (Figure 4 A2 top). Individual traces from two PY neurons (one from each cluster) show spontaneous background activity in both clusters (Figure 4 A2 bottom). The mean Na^+^/K^+^ pump current within each cluster revealed synchronized resting-state fluctuations (Figure 4B). Fourier transforms of the mean Na^+^/K^+^ pump currents from both clusters revealed a power spectrum peak around 0.02 Hz (Figure 4C left). Importantly, fluctuations of the mean membrane potential and mean Na^+^/K^+^ current of both clusters revealed positive cross correlation (Figure 4C middle & right respectively).

**Figure 4.**
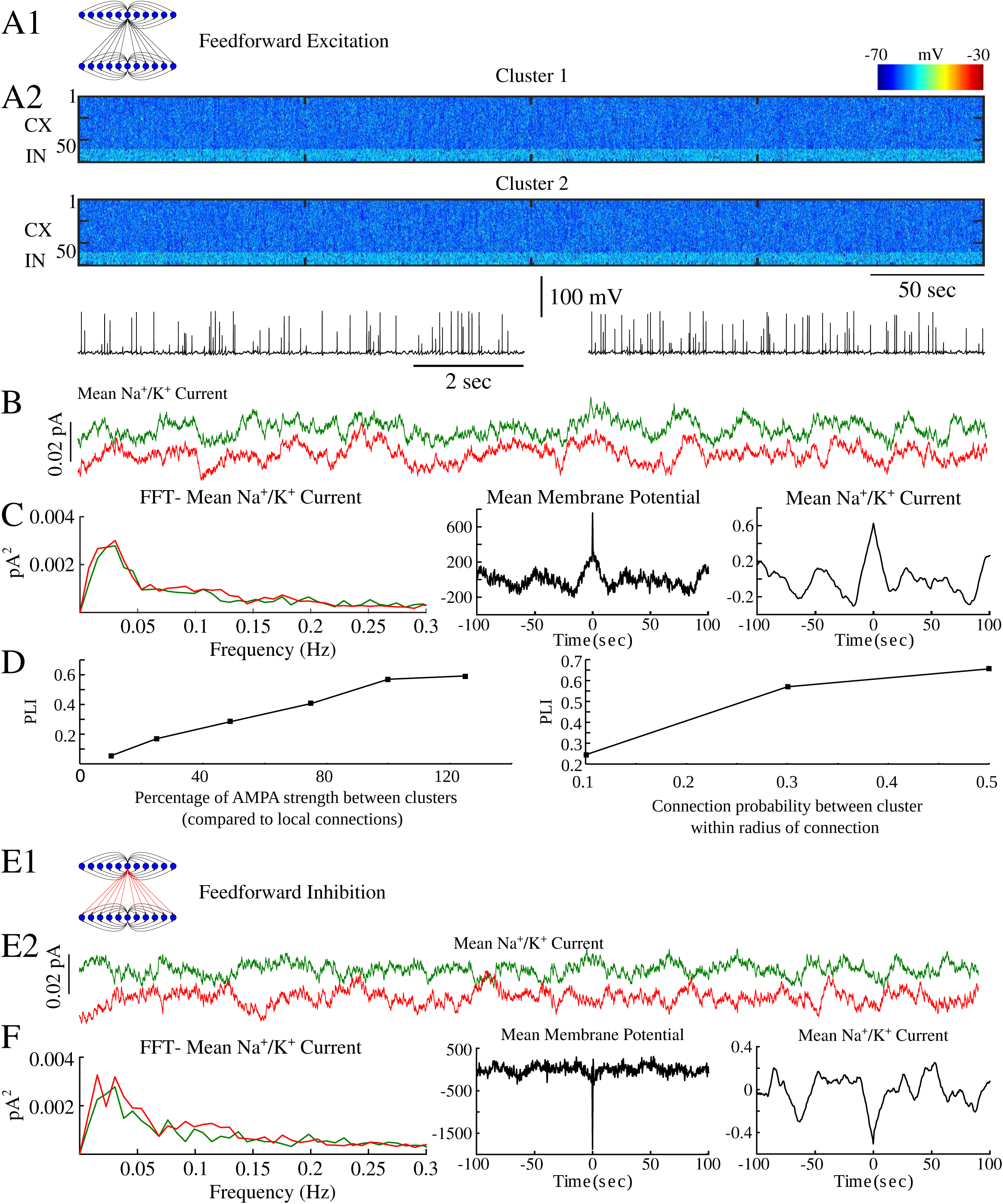
Longe-range projections synchronize resting-state fluctuations. **A1,** Cartoon of feedforward excitation. **A2,** Spontaneous activity from both neuron clusters connected by feedforward excitation. **B,** Mean Na^+^/K^+^ pump currents for each cluster in A. **C,** FFTs of Na^+^/K^+^ pump currents in B (left), cross-correlation of mean filtered membrane potentials from clusters in A (middle), and cross-correlation of mean Na^+^/K^+^ pump currents from B (right). **D,** Phase-locking index (PLI) as a function of feedforward AMPA strength (left) and connection probability (right). **E1,** Cartoon of feedforward inhibition. **E2,** Mean Na^+^/K^+^ pump currents for each cluster connected through feedforward inhibition. **F,** FFTs of Na^+^/K^+^ pump currents in E2 (left), cross-correlation of mean filtered membrane potentials from clusters with feedforward inhibition (middle), and cross-correlation of mean Na^+^/K^+^ pump currents from E2 (right).

We next varied the AMPA connection strength of the long-range connections between the two clusters to test its effect on the synchronization between the resting fluctuations of the two clusters. We computed the phase-locking index (PLI) between the mean Na^+^/K^+^ pump currents of the two clusters for different AMPA strengths. Increasing the AMPA strength of the long-range connections resulted in the higher phase-locking between the two signals (Figure 4D left). Similarly, increasing connection probability between the two clusters resulted in an increase of the phase-locking (Figure 4D right). Importantly, even for relatively low connection strength and probability, the PLI remained significantly higher than that for two completely disconnected clusters.

In vivo, in addition to the positively correlated resting-state fluctuations in distinct regions, negatively or anti-correlated resting-state fluctuations have been observed (38-40). We found that increasing the strength of the AMPA connection between excitatory PY neurons of the one cluster and the inhibitory IN neurons of another cluster resulted in the negatively correlated resting-state fluctuations (Figure 4E). This negatively correlated or anti-phasic activity was observed in both the filtered mean membrane potential and the mean Na^+^/K^+^ pump currents of the clusters. Cross correlation analysis of these two signals revealed a clear negative correlation between the resting-state fluctuations, as shown by the negative peaks in figure 4F (middle & right). Together, these findings suggest that the mechanisms proposed in our model can account for individual cluster resting-state fluctuations as well as for positive and negatively correlated fluctuations between distinct brain regions.

Finally, to test whether our model could explain *in vivo* data that revealed correlations between structural and functional connectivity in the Macaque brain (8, 11, 25, 26), we modeled 58 different brain regions of the Macaque brain using connectivity information gathered from the CoCoMac structural connectivity database (http://cocomac.g-node.org). Each of the 58 regions was modeled as a cluster of 50 excitatory neurons and 10 inhibitory interneurons with connectivity within a cluster similar to that shown in Figure 1 A1. Long-range excitatory PY-PY connections were formed between clusters based on the CoCoMac structural connectivity data set (http://cocomac.g-node.org). We computed the correlation coefficient between Na^+^/K^+^ pump currents in different clusters for every possible pair of clusters. This analysis revealed the groups of clusters which showed high degree of correlation (Figure 5A, middle panel). To quantify the relationship between structural and functional connectivity, we next computed the correlation coefficient between clusters (brain regions) showing significant correlation of the mean Na^+^/K^+^ pump currents (Figure 5A, middle) and the clusters with strong structural connectivity (Figure 5A, left panel). We found a significant correlation between the functional and structural connectivity (r = 0.20508) (Figure 5A, right panel). Thus, we concluded that a network with local (cluster specific) ion concentration dynamics and long-range synaptic connectivity between clusters can account for the experimentally observed relationship between structural and functional connectivity. Recent data revealed the dynamic nature of functional connectivity (41-44). It has been shown that the strength of the function connectivity computed from the resting-state fluctuations between regions comprising the default mode network was varying in time (3). To check for a similar characteristic in our model, we computed the correlation coefficients for ten subsequent 60sec bins (Figure 5B). Similar to experimental findings, our model revealed dynamic changes of the functional connectivity (Figure 5B). The network exhibited both instances of strong and weak functional connectivity between specific clusters at different time epochs. In particular, regions that showed initially weak functional connectivity developed strong connectivity before returning to a low connectivity state.

**Figure 5.**
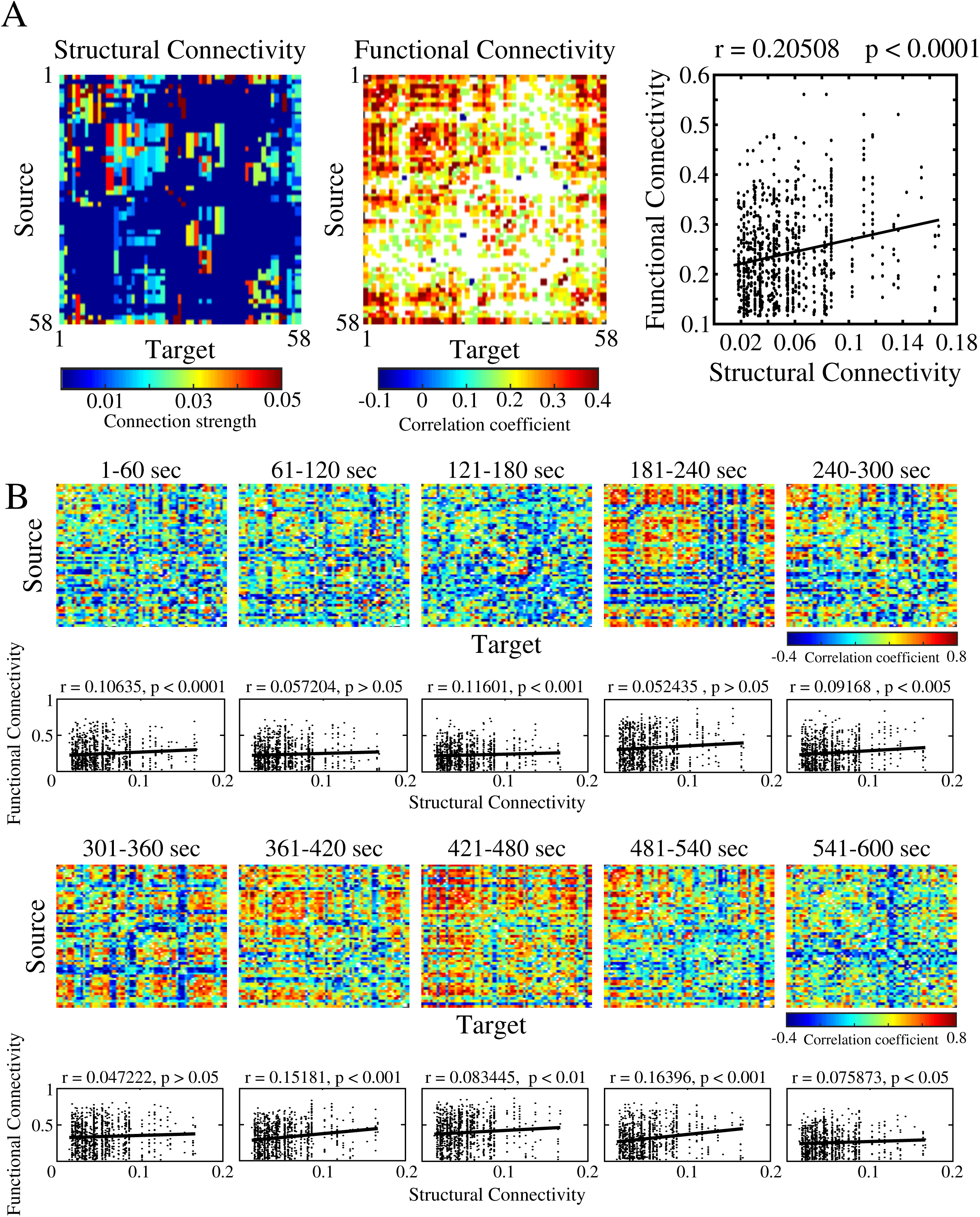
Macaque simulations. **A,** Left, structural connectivity matrix for macaque network. Middle, computed functional connectivity showing only significant correlations (Bonferroni corrected for multiple comparisons). Right, correlation of functional and structural connectivity. **B,** Heatmaps, function connectivity computed for consecutive 60sec time windows. Bottom plots, correlations between functional connectivity computed in corresponding heatmap and structural connectivity.

## Discussion

In this study, we tested hypothesis that dynamics of the ion concentrations, regulated through the neuronal and glial activity, form the basis of the resting-state fluctuations in the brain. Comprising only about 2% of the total body weight of an average adult human, the brain is responsible for up to 20% of the total energy consumption (18). Task-evoked responses generally increase brain energy consumption by less than 5% (45). Though so much energy is consumed in order to maintain a baseline level of activity, little is known about its use, including spontaneous resting-state activity in the brain. It was first observed by Biswal et al. 1995 that the spontaneous background fluctuations recorded during fMRI scans were coherent between functionally related brain regions (19). Since then, additional studies have shown similar coherent resting-state activity between regions comprising functional networks such as the default mode network and executive control network (1, 2, 4, 6-8, 13, 18, 45, 46). Interestingly, infra-slow (<0.2Hz) resting-state fluctuations have been observed in various cognitive states (4, 7, 13, 17), and can exhibit modified temporal coherence patterns in various neurological and psychiatric disorders (1, 2, 6, 20, 21). In our new study we proposed and tested the hypothesis that resting-state fluctuations may depend on ion concentration dynamics, specifically [K^+^]_o_ fluctuations, and that the phase coherence of the infra-slow activities between distinct brain regions depends on the long-range synaptic connectivity between these regions. Our model based on the CoCoMac structural connectivity database explained the relationship between structural and functional connectivity revealed in the studies of the resting state activity in the Macaque brain.

The characteristic period of resting-state fluctuations is on the order of 50-100sec. Very few neural processes are known to act at such slow time scale. In our new study, we demonstrate that the ion concentrations may spontaneously vary with a very slow time scale and could act as the modulator of the neural activity leading to emergence of resting-state fluctuations. The underlying mechanisms generating these infra-slow fluctuations may rest in the slow processes leading to the gradual accumulation and removal of the extracellular K^+^. The positive-feedback loop between extracellular K^+^ concentration and neuronal firing rate may lead to progressive increase in the firing rate and extracellular K^+^. Due to its non-linear nature, the Na^+^/K^+^ pump is significantly more activated as the extracellular K^+^ reaches a critical value controlled by the pump half-activation threshold, which then results in the reduction of the extracellular K^+^ and the neuronal firing rate. Indeed, experimental data suggested that the ion concentrations may have slow dynamics similar to the time course of the resting-state fluctuations (27-29, 32-36, 47). Our previous modeling studies revealed that the ion concentration dynamics may lead to the slow (<0.2 Hz) quasi-periodic transitions between distinct network states (bursting and tonic firing) (27-30, 32-36, 47, 48). Other studies have suggested the role of slow processes involving ion dynamics and Na^+^/K^+^ pump activity in information processing (49, 50).

The gradual accumulation of [K^+^]_o_ has been suggested to initiate a positive-feedback loop resulting the heightened neuronal excitability and development of seizure discharges (28-30, 34, 37). Studies in patients with epilepsy revealed abnormal resting-state fluctuations (20, 22, 23). Indeed, it has been demonstrated that the amplitude of resting-state fluctuations in epileptic patients is increased as compared to the healthy individuals (22). Interestingly, our previous work demonstrated that homeostatic up-regulation of excitatory connections, following trauma, may lead to rewiring long-range cortical connectivity (37). Taken together, it may explain differences in the resting-state fluctuation properties in epileptic patients and healthy individuals.

Studies in animals revealed a correlation between anatomical structure and functional connectivity (8, 11, 25, 26). It was proposed with computer models that the time scale of infra-slow fluctuations could be a result of transient bouts of synchrony between clusters of nodes, and that the functional connectivity arising in the network strongly reflected the underlying structural architecture (11, 25). However, these earlier modeling studies were based on the phenomenological mean field type models. Here we apply biophysically realistic network model of the cortical circuit implementing populations of excitatory and inhibitory cells and the ion concentration dynamics to explain infra-slow activity. In agreement with previous data (11, 25), functional connectivity in our model, computed over a long time window, reliably reflected the underlying structural connectivity.

Experimental work has shown that the lack of monosynaptic connections between brain regions does not accurately predict the absence of functional connectivity between those regions (7). However, coupling strength between neuron clusters has been suggested to influence the strength of correlated and anti-correlated activity between nodes (25). In our model, the correlation and phase-locking of infra-slow fluctuations between clusters of neurons reflected the strength of excitatory connections between clusters (figure 4D). Stronger excitatory projections resulted in correlated activity and phase-locking of the infra-slow fluctuations. We found that feedforward inhibition could give rise to anti-correlated activity between clusters (figure 4E and F) and we observed dynamic changes of the functional connectivity, in agreement to the experimental studies (41-44).

Recent experimental work has shown that fluctuations in the fMRI BOLD signals are well correlated with the local field potentials (LFPs) (9, 10, 12, 15-17, 24, 51). Gamma band activity and slow cortical potentials (SCPs) have been shown to correlate with BOLD signals (12, 18, 24). Similar to BOLD signals, SCPs show coherent activity patterns across various cognitive states. It has been suggested that the time scale of these SCPs may reflect slow metabolic processes occurring within populations of neurons (18, 24). Aerobic glycolysis accounts for ~15% of the glucose metabolized at rest (24). The main glucose metabolite is ATP, most of which is consumed by the Na^+^/K^+^ pump in order to maintain ionic gradients (52, 53). Modifying glucose concentration has shown to modulate slow wave oscillations *in vitro* in rat entorhinal cortex (52). In agreement with these results, our model suggests that Na^+^/K^+^ pump activity may influence the properties of the infra-slow fluctuations. Na^+^/K^+^ pump activity is reflected in fMRI BOLD signals as oxygenated blood flow increases to brain regions exhibiting increased neuronal activity (54). It suggests that taking into account effects of the ion concentrations and Na^+^/K^+^ pump dynamics, as implemented in our model, may be necessary to accurately capture the biophysical mechanisms leading to generation of infra-slow fluctuations in fMRI recordings.

### Methods and Materials

*Intrinsic properties and ion dynamics of excitatory and inhibitory neuron models*. Our network model with dynamics ion concentrations has been described in detail elsewhere (27, 28, 30, 31, 37). Briefly, both excitatory pyramidal cells (PYs) and inhibitory interneurons (INs) were modeled as two-compartment neurons with an axosomatic and a dendritic compartment. The evolution of voltage in time for each compartment was described by the following equations:

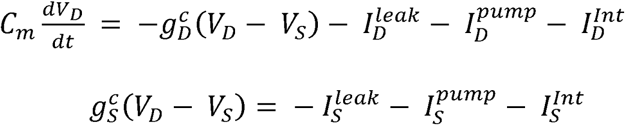

where *V*_*D,S*_ are the voltages of the dendritic and axosomatic compartments (respectively), 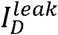 and 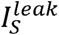 are the sum of the ionic leak currents, 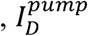 and 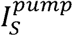 are the sum of the Na^+^ and K^+^ currents through the Na^+^/K^+^ pump, and 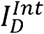 and 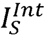 are the intrinsic currents for the dendritic and axosomatic compartments respectively. Each of these compartments contained conductance-based Hodgkin-Huxley type ionic currents. The ionic current intrinsic to the dendritic compartment include the fast sodium (*I*_*Na*_), persistent sodium current (*I*_*NaP*_), slowly activating potassium current (*I*_*Km*_), high-threshold calcium current (*I*_*Ca*_), calcium-activated potassium current (*I*_*KCa*_) hyperpolarization-activated depolarizing mix cationic currents (*I*_*h*_), and leak currents (30, 31, 37). The axosomatic compartment contains the fast sodium current (*I*_*Na*_), the persistent sodium current (*I*_*KaP*_), delayed-rectifier potassium current (*I*_*Kv*_), and the sodium-activated potassium current (*I*_*KNa*_). Na^+^/K^+^ pump Na^+^ and K^+^ regulation and KCC2 cotransporter Cl^-^ extrusion were included in both neuron types. Additionally, dynamic ion concentrations for extracellular and intracellular Na^+^ and K^+^ as well as intracellular Cl^-^ and Ca^2^+ were determined by the intrinsic currents, transporter-mediated currents, leak currents, extracellular and intracellular diffusion, and glial [K^+^]_o_ buffering as described in the following equations:

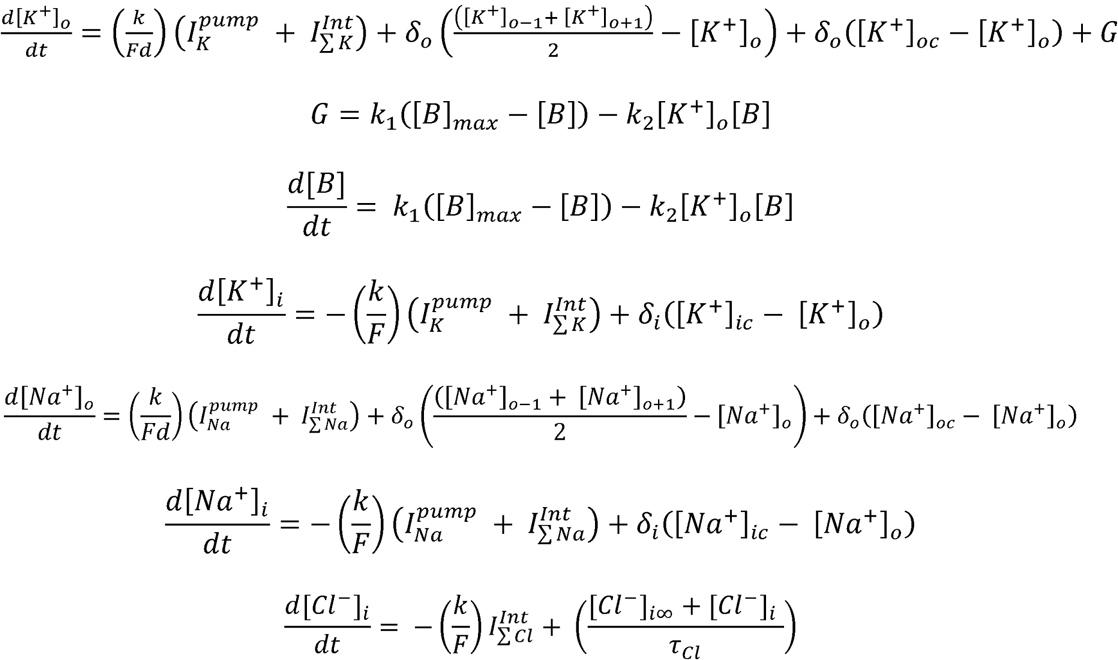

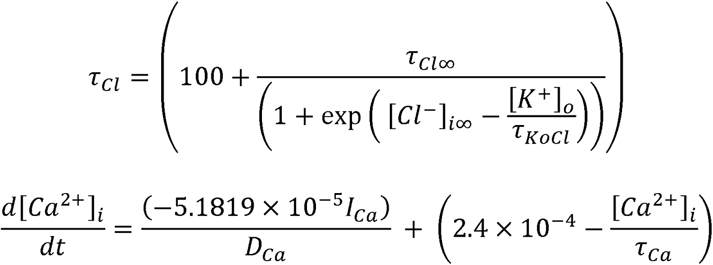

where *F* = 96489 C/mol, *d* = 0.15 is the ratio of the extracellular compartment volume to surface area, the conversion factor *k* = 10, *δ*_*o*_ is the diffusion coefficient, [*K*^+^]_*oc*_ and [*Na*^+^]_*oc*_ are the K^+^ and Na^+^ concentrations in the adjacent compartments, and [*K*^+^]_o-1_, *K*^+^]_o+1_, *Na*^+^]_o-1_, and [*Na*^+^]_*o*+1_ are the concentrations of K^+^ and Na^+^ in neighboring cells respectively. Glial buffering of extracellular K^+^ (*G*) was modeled as a free buffer ([*B*]) with total buffer ([*B*]_*max*_) = 500mM. The [*B*] K^+^ binding and unbinding rates (*k*_1_ and *r*_2_ respectively) were given by *k*_1_ = 0.0008 and *k*_2_ = *k*_1_(1 + *exp*(([*K*^+^]_o_ – [*K*^+^]_*oth*_)/(‒1.05))), where [*K*^+^]_*oth*_ = 15mM is the half activation concentration of [*K*^+^]_*o*_. [*Cl*^-^]_*i∞*_ = 5mM, *τ*_*Cl*∞_ = 2x10^4^, and *τ*_*KoCl*_ = 0.08s. *τ*_*Ca*_ and *D*_*Ca*_ were set to 300ms and 0.85 respectively. Extracellular K^+^ was also allowed to diffuse between the two compartments as well as between neighboring cells of the same type (ie. diffusion between PY-PYs and IN-INs).

*Network and synaptic properties*. This network model was modeled as a one dimensional network and consisted of 50 excitatory PY neurons and 10 inhibitory IN neurons (unless otherwise specified). Each PY neuron made local excitatory connections onto 10 other PY neurons and received 10 excitatory connections from other PY neurons. PY neurons also formed excitatory connections onto inhibitory IN neurons. Each PY projected onto one IN and each IN formed inhibitory connections onto 5 PY neurons. Excitatory connections were mediated by AMPA and NMDA conductances (11 nS and 1 nS, respectively), and inhibitory connections were mediated by GABA_A_ conductances (11 nS) such as those described previously (30, 31, 37). Excitatory connections from PY neurons onto IN neurons were mediated by AMPA and NMDA conductances (3.5 nS and 0.35 nS, respectively). To model *in vivo* conditions, all neurons of both types received additional afferent excitatory input as a random Poisson process.

*Single and double cluster network connectivity*. For single cluster networks, a 1:5 ratio of IN to PY neurons was kept, regardless of the network size. Local connections within the cluster were formed with twenty percent connection probability (unless otherwise stated). As previously mentioned, local diffusion of K^+^ was allowed between PY-PY and IN-IN but not between PY and IN neurons.

To study the synchrony between clusters of neurons, we generated networks consisting of two clusters of neurons. These networks consisted on 100 neurons, with two clusters of 50 neurons. Similar to the above mentioned neuron cluster properties, 1:5 ratio of IN to PY neurons was kept for each cluster. Additionally, ion diffusion was not allowed between the clusters. In order to connect the two clusters, PY neurons were allowed to make long-range connections from one cluster to the other. These long-range connections were PY-PY and PY-IN connections with the same connection radius as described for local connections within a cluster. Connection probabilities and connection strengths for these long-range connections were varied in ordered to study how these connection properties influence correlation between cluster oscillations.

*Phase-locking analysis*. Phase-locking was used to study the synchrony between two fluctuations in two clusters. The phase-locking index (PLI) between two clusters (*x*_1_ and *x*_1_) was computed as follows:

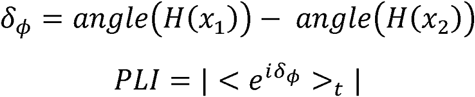

where *H* is the Hilbert transform, and *δ*_*ϕ*_ is the difference in angle between the two time series. Both time series were band-pass filtered between 0.001 and 0.1 Hz before the Hilbert transform was applied.

*Macaque simulations*. We implemented the structural connectivity of 58 macaque brain regions. Connection strengths between brain regions were extracted from the CoCoMac database (http://cocomac.g-node.org). Functional connectivity was computed as the correlation coefficients between mean Na^+^/K^+^ pump currents from individual clusters. Significance values were Bonferroni corrected to correct for multiple comparisons. For analysis of the dynamic nature of the resting-state fluctuations, 60sec time windows were used to compute correlation coefficients. An agglomerative hierarchical clustering (using ward distance metric) was applied to the correlation coefficient matrix consisting of all combination of clusters and was used for visualization used in Fig 5.

## Acknowledgements

This study was supported by Office of Naval Research Multidisciplinary University Research Initiative Grant N000141612829. Oscar C. González was supported by the NSF Graduate Research Fellowship under grant DGE-1326120.

